# X-ray mediated scintillation increases synaptic activity via Cerium-doped LSO and Channelrhodopsin-2

**DOI:** 10.1101/2020.08.29.273359

**Authors:** Aundrea F. Bartley, Máté Fischer, Micah E. Bagley, Justin A. Barnes, Mary K. Burdette, Kelli E. Cannon, Mark S. Bolding, Stephen H. Foulger, Lori L. McMahon, Jason P. Weick, Lynn E. Dobrunz

## Abstract

Optogenetics is a widely used tool for studying neural circuits. However, non-invasive methods for light delivery in the brain are needed to avoid physical damage typically caused by intracranial insertion of light guides. An innovative strategy could employ X-ray activation of radioluminescent particles (RLPs) to emit localized light. We previously reported that RLPs composed of cerium doped lutetium oxyorthosilicate (LSO:Ce), an inorganic scintillator that emits blue light, are biocompatible with neuronal function and synaptic transmission. However, little is known about the consequences of acute X-ray exposure on synaptic function and long-term plasticity. Furthermore, modulation of neuronal or synaptic function by X-ray induced radioluminescence from RLPs has not yet been demonstrated. Here we show that 30 minutes of X-ray exposure at a rate of 0.042 Gy/second caused no change in the strength of basal glutamatergic transmission during extracellular dendritic field recordings in mouse hippocampal slices. Additionally, long-term potentiation (LTP), a robust measure of synaptic integrity, was able to be induced after X-ray exposure and expressed at a magnitude not different from control conditions (absence of X-rays). This is important as synaptic plasticity is critical to learning and memory. Next, we used molecular and electrophysiological approaches to determine if X-ray dependent radioluminescence emitted from RLPs can activate light sensitive proteins. We found that X-ray stimulation of RLPs elevated cAMP levels in HEK293T cells expressing OptoXR, a chimeric opsin receptor that combines the extracellular light-sensitive domain of channelrhodopsin-2 (ChR2) with an intracellular second messenger signaling cascade. This demonstrates that X-ray radioluminescence from LSO:Ce particles can activate OptoXR. Next, we tested whether X-ray activation of the RLPs can enhance synaptic activity in whole-cell recordings from hippocampal neurons expressing ChR2, both in cell culture and acute hippocampal slices. Importantly, X-ray radioluminescence caused an increase in the frequency of spontaneous excitatory postsynaptic currents (sEPSCs) in both systems, indicating activation of ChR2 and excitation of neurons. Together, our results show that X-ray activation of LSO:Ce particles can heighten cellular and synaptic function. The combination of LSO:Ce inorganic scintillators and X-rays is therefore a viable method for optogenetics as an alternative to more invasive light delivery methods.

## Introduction

Optogenetics is a tool that has allowed the neuroscience community to expand our understanding about individual neuronal cells and specific brain circuits during behavior and in various disease states (Gradinaru et al., 2009; Yizhar et al., 2011; Lim et al., 2013; Gunaydin et al., 2014; Emiliani et al., 2015; Fenno et al., 2015; Rost et al., 2017; Selimbeyoglu et al., 2017; Barnett et al., 2018). The use of various types of light activated ion channels is a fundamental aspect of optogenetics to either cause membrane potential depolarization or hyperpolarization to induce or suppress action potential firing, respectively. As with all techniques, improvements to the method are constantly being developed (Rein and Deussing, 2012; Lim et al., 2013; Lin et al., 2017; Chen et al., 2018) and there is a need for less invasive methods of light delivery to the brain in vivo.. The most common approach requires implantation of fiber optic waveguides (fibers) or light emitting diodes (LEDs) which are several hundred microns in size and cause damage to delicate brain tissues and multiple brain regions when implanted into deep brain structures (Aravanis et al., 2007; Ozden et al., 2013; Canales et al., 2018). In order to activate the large population of cells needed to modulate behavior, the placement of light emitting devices can be far from the region of interest. These limitations combined with the fact that brain tissue absorbs and scatters light requires relatively high light intensities to be produced by the light source. This enhanced illumination can cause increases in local temperatures (Senova et al., 2017) with the consequence of increasing neuronal firing rates (Stujenske et al., 2015). Clearly, technology advances are needed to provide less invasive options for light delivery for synaptic circuit control for in vivo optogenetics (Chen et al., 2018; Matsubara et al., 2019).

One potential strategy for less invasive light delivery to the brain is the use of X-rays to activate radioluminescent materials (Shuba, 2014; French et al., 2018; Bartley et al., 2019; Matsubara et al., 2019). Radioluminescence is produced by exposing scintillating material to ionizing radiation, thereby causing generation of visible light. Radioluminescent particles (RLPs) are developed from inorganic scintillator material that can emit light at various wavelengths based upon its composition. The use of RLPs would allow the light to be generated locally, overcoming the issue of light attenuation and the need for high power intensities. A common inorganic scintillator is Cerium-doped lutetium oxyorthosilicate (LSO:Ce). LSO:Ce material has a high light output (Melcher and Schweitzer, 1991; Roy et al., 2013), and its X-ray activation has been shown to emit wavelengths of light needed to activate channelrhodopsin-2 (ChR2) (Melcher and Schweitzer, 1991). Previously, we showed that LSO:Ce particles had minimal effects on neuronal function and synaptic transmission (Bartley et al., 2019). Together, this makes it an ideal material to determine if radioluminescence from RLPs generated by X-ray exposure can activate ChR2 to modulate synaptic circuits.

X-rays are able to penetrate the skull and reach deep brain structures without causing damage. As a result, X-rays applied from outside the skull could be used to activate RLPs in the brain, thus locally producing light needed to activate light-sensitive effectors such as ChR2. This would remove the need for implanted LEDs, reducing physical damage to brain tissue. However, X-irradiation can impair neurogenesis with prolonged exposure (Mizumatsu et al., 2003; Cacao et al., 2018). Little is known about how X-ray exposure affects synaptic transmission and circuit function, especially with acute applications. At the neuromuscular synapse, a high dose of X-rays has been shown to diminish synaptic transmission (Portela et al., 1977). Additionally, a high dose of X-rays causes impairment in memory formation that is not only due to alterations with neurogenesis (Puspitasari et al., 2016). But, the level of exposure that would be needed for a non-invasive optogenetic technique to work and its effect on circuit function is unknown.

Here we demonstrate that acute exposure of X-rays (<80 Gy) had no effect on basal synaptic excitatory field potentials (fEPSPs) at hippocampal CA3-CA1 synapses. The use of X-irradiation did not impair induction or expression of long-term potentiation (LTP). Importantly, we demonstrate that light emitted from LSO:Ce microparticles following X-ray exposure is able to activate ChR2 and modulate cellular and synaptic function. Together, these results show that the combination of X-rays and LSO:Ce particles are neither toxic to brain slices nor detrimental to synaptic function. Therefore, the use of RLPs and X-rays is a potential method for a less invasive form of light delivery to the brain for in vivo optogenetics.

## Materials and Methods

### 1. Animals

Approval was obtained for all experimental protocols from the University of Alabama at Birmingham and University of New Mexico Institutional Animal Care and Use Committees. All experiments were conducted in accordance with the Guide for the Care and Use of Laboratory Animals adopted by the National Institutes of Health. C57Bl/6J mice were housed in a 22 ± 2° C room with food and water ad libitum. Four-week-old C57Bl/6J mice were purchased from JAX Labs (JAX# 000664) for slice electrophysiology experiments. For a subset of experiments, adult Emx:ChR2 mice were used. These mice had expression of channelrhodopsin-2 (ChR2) in excitatory neurons, which was accomplished by crossing Emx-cre mice (B6.129S2-Emx1^tm1(cre)Krj^/J, JAX# 005628) (Gorski et al., 2002) with floxed ChR2/EYFP mice (Ai32 (RCL-ChR2 (H134R)/EYFP, JAX # 012569) (Madisen et al., 2012). Emx:ChR2 mice were housed in a 26 ± 2°C room with food and water ad libitum. Mice were housed with the whole litter until weaned (P23). Weaned mice were housed with no more than 7 mice in a cage. Mouse genotypes were determined from tail biopsies using real time PCR with specific probes designed for cre and EYFP (Transnetyx, Cordova, TN).

### 2. Primary Hippocampal Neuronal Cultures

Hippocampal cultures were made using previously established methods (Chander et al., 2019). Briefly, brains from P0-P1 C57Bl/6J pups (The Jackson Laboratory, Bar Harbor, ME) were isolated, and the hippocampus was dissected in ice-cold HBSS solution (Sigma, St. Louis, MO) supplemented with 20% FBS and NaHCO3 (4.2mM), HEPES (1mM; Sigma), pH 7.4. Dissected hippocampi were digested for 10 min with 0.25% Trypsin (Thermo Fisher Scientific), then washed and dissociated using fire polished Pasteur pipettes of decreasing diameter in ice cold HBSS containing DNase (1500 U; Sigma). The cells were pelleted, resuspended in plating media and plated at a density of 4-5×105 cells/12-mm coverslip (Electron Microscopy Sciences, Hatfield, PA) coated with poly-Ornithine (0.1mg/ml; Sigma; Cat. #4638) and laminin (5μg/ml; Thermo Fisher Scientific). Cells were allowed to adhere for 15 min before addition of 0.5ml of plating media containing Neurobasal supplemented with 1X B27, 2mM Glutamax, 0.5mg/ml Pen/Strep and 5% FBS (all from Thermo Fisher Scientific) for the first 24h. Half of the media was removed and replaced with serum-free media after 24h. Half of the media was removed and replaced after 48h supplemented and with 4μM cytosine 1-β-d-arabinofuranoside (Ara-C; Sigma). Neurons were fed by replacing half the volume of spent media with fresh media without serum or Ara-C every week thereafter.

### 3. Viral Production and Transduction

Lentivirus was produced in HEK293T cells using calcium phosphate transfection of a 15 μg total mixture of lentiviral targeting vector (Syn-ChR2(H134R)-mCherry; (Weick et al., 2011) referred to here as ChR2-mCherry) or EF1a-DIO Opto β2-AR-EYFP; (Airan et al., 2009), referred to here as OptoXR-EYFP) and packaging plasmids psPax2 and pMD2.G (Addgene, Cat. # 12260 and Cat. # 12259 respectively) at a ratio of 3:2:1 for lentivirus production. 36-48 h post transfection lentivirus containing media was harvested and concentrated using Lenti-X concentrator (Takara, Mountain View, CA) as per manufacturer’s recommendations. Viral pellets were resuspended in ice cold DMEM, aliquoted, and frozen at −80°C until use, showing a viral titer of >10^6^ IFU/ml as measured by Lenti-X GoStix (Takara, Mountain View, CA). To develop stably expressing OptoXR-EYFP cells, HEK293T cell cultures were transduced with OptoXR-EYFP using a multiplicity of infection (MOI) of 1. After allowing 2-3 days for the expression of the lentiviral products, HEK293T cells were dissociated and sorted via flow cytometry into EYFP-positive and −negative subpopulations according to previously published methods (Basu et al., 2010). Following cell sorting, EYFP^+^ cells were plated in 10cm dish format and propagated for subsequent assay. Neuronal cultures were transduced with ChR2-mCherry using a MOI of 1 at 5-7 days in vitro (DIV).

### 4. X-Ray Exposure

For electrophysiology experiments and experiments using different opsins, X-ray exposure was generated by a Mini-X silver target X-ray unit (Amptek, Bedford, MA) with both the nozzle and collimator removed. The unit was operated at 50 kV and 79 μA. The distance between the sample and the actual X-ray source was approximately 3 to 3.5 cm. The output (C) of the Mini-X was measured with a RadCal 9010 X-ray dosimeter in a RadCal 10 x 6 ionization chamber. Using the equation below, the radiation dose with the X-ray source approximately 3.5 cm from the neurons provides a radiation dose rate of 0.042 Gy/s, which is relatively close to our estimates with the dosimeter stickers.

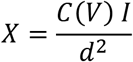

where X is the radiation dose rate, I is the tube current, d is the distance from the X-ray focal spot, and C is a function of tube voltage (V), which was measured at 50 kV at a distance of 7 cm. Additionally, RADSticker dosimeter stickers (JP Laboratories, Inc.) were used to approximate the radiation dose on the electrophysiology setups, and was approximately 0.05 Gy/s and is very similar to our calculations.

X-rays were applied for 30 minutes before the application of the high frequency stimulation (HFS) protocol used to induce LTP. For activation of ChR2 experiments in primary neuronal cultures, spontaneous excitatory postsynaptic currents (sEPSCs) were recorded for 30 seconds without X-rays and then x-irradiation was applied for 30 seconds, this cycle was repeated 3 times. For activation of ChR2 experiments, in acute hippocampal slices, the pattern was 2 minutes of recording without x-irradiation followed by 2 minutes with X-rays and was repeated 3 times.

The X-ray radioluminescence spectra of the LSO:Ce microparticles was obtained using a fiber bundle (Oriel) connected to a MicroHR monochromator (Horiba Jobin Yvon) equipped with a CCD detector (Synapse). The sample was irradiated with a mini X-ray tube (Amptek Inc., MA, USA) equipped with a silver target operating at a tube voltage of 50 kV and a tube current of 79 μA. The signal was collected on a grating with 300 line mm−1 and a blaze of 600 nm with an exposure time of 1.2 seconds.

### 5. Whole Cell Recordings from Cultured Neurons

Whole cell voltage clamp recordings of cultured hippocampal neurons were performed as previously described (Chander et al., 2019) with minor modifications. The extracellular solution was a modified Hanks’ balanced salt solution (HBSS) that contained (in mM): 140 NaCl, 3 KCl, 2 CaCl2, 1 MgCl2, 15 HEPES, and 23 glucose, pH 7.4, 300 mOsm. Recording pipettes with resistances of 3 to 5 MΩ were filled with an intracellular recording solution containing the following (in mM): 121 K-gluconate, 20 KCl, 2 MgCl2, 10 EGTA, 10 HEPES acid, 2 Mg2+-ATP, 0.2 Na+-GTP, pH 7.2, 290 mOsm. Neurons were visualized using an Olympus Optical BX51WI microscope (Olympus Corp., Tokyo, Japan) with differential interference contrast optics at 40x. Recordings were obtained using a MultiClamp 700B amplifier (Molecular Devices, Sunnyvale, CA), filtered at 4 kHz and sampled at 100 kHz using a Digidata 1322A analog-to-digital converter (Molecular Devices). Whole-cell capacitance was fully compensated but series resistance was not compensated. Access resistance was monitored before and after recordings, and cells with resistances greater than 20 MΩ at either point were discarded from analyses. Spontaneous excitatory postsynaptic currents (sEPSCs) were recorded in the absence of tetrodotoxin but Picrotoxin (50 μM; Tocris, Bristol, UK) was bath applied in the external solution to isolate excitatory currents. sEPSCs were measured using a holding potential of −70mV and all recordings were performed at 32°C. sEPSCs were analyzed using MiniAnalysis software (Synaptosoft, Fort Lee, NJ). 10μl of LSO:Ce doped RLP solution was applied to coverslips containing cultured neurons immediately prior to recording. As the particles do not readily wash off, the final concentration of LSO:Ce RLPs was between 0.1 to 0.5 mg/mL. The irradiation protocol was applied two minutes after start of whole-cell recording (break in), to allow perfusion with the internal solution.

### 6. Electrophysiology in Acute Hippocampal Slices

#### 6.1 Mouse Hippocampal Slice Preparation

Young adult male C57Bl/6J mice (age 4 weeks; JAX# 000664) were anesthetized with isoflurane, decapitated, and brains removed; 400 μm coronal slices from dorsal hippocampus were made on a VT1000P vibratome (Leica Biosystems) in oxygenated (95%O2/5%CO2) ice-cold high sucrose cutting solution (in mm as follows: 85.0 NaCl, 2.5 KCl, 4.0 MgSO_4_, 0.5 CaCl_2_, 1.25 NaH2PO_4_, 25.0 glucose, 75.0 sucrose). After cutting, slices were held at room temperature with continuously oxygenated standard artificial cerebral spinal fluid (aCSF) (in mm as follows: 119.0 NaCl, 2.5 KCl, 1.3 MgSO_4_, 2.5 CaCl_2_, 1.0 NaH_2_PO_4_, 26.0 NaHCO_3_, 11.0 glucose).

Male and female Emx:ChR2 mice, 3 to 7 months of age, were anesthetized with isoflurane and sacrificed by decapitation using a rodent guillotine. The brains were rapidly removed and placed in ice cold dissection solution containing the following (in mM): 135 N-Methyl-D-glucamine, 1.5 KCl, 1.5 KH_2_PO_4_, 0. 5 CaCl2, 3.5 MgCl2, 23 NaHCO3, 0.4 L-Ascorbic acid, and 10 glucose, bubbled with 95% O2/5% CO2, pH 7.35– 7.45, and osmolarity 295–305 (Albertson et al., 2017). This dissection solution has been shown to improve slice health for older age animals (Ting et al., 2018). A vibratome (Campden 7000smz-2, Lafayette Instrument) was used to cut 300 μM thick hippocampal brain slices. The slices were maintained in oxygenated recovery solution containing (in mM) 120 NaCl, 3.5 KCl, 0.7 CaCl_2_, 4.0 MgCl_2_, 1.25 NaH_2_PO_4_, 26 NaHCO_3_, and 10 glucose and kept at room temperature. Slices were stored at room temperature in a humidified oxygenated interface recovery chamber in 2 mLs of solution on top of a piece of filter paper and bubbled with 95% O2/5% CO2 for > 1.5 hours before recording.

#### 6.2 Incubation of LSO:Ce microparticles with Acute Hippocampal Slices

Acute hippocampal slices were allowed to recover for an hour before application of either vehicle (artificial cerebral spinal fluid, aCSF) or 0.125 to 0.5 mg/mL of LSO:Ce radioluminescent microparticles. This is an underestimate of the concentration of the particles on the slice as they do not evenly distribute in the solution. The slices were incubated with and without LSO:Ce microparticles for at least 30 minutes.

#### 6.3 Electrophysiology - Field Recordings

Slices were interleaved between with and without X-ray exposure to control for possible changes in slice health during the recordings. Extracellular field excitatory postsynaptic potentials (fEPSPs) were recorded from the dendritic region in hippocampal area CA1 using a submersion chamber perfused with standard aCSF at 24-25° C. All data were obtained using the electrophysiology data acquisition software pClamp10 (Molecular Devices, LLC, Sunnyvale, CA.) and analyzed using Clampfit within the pClamp10 suite, and Graphpad Prism 7 (GraphPad Software, Inc.). For CA3-CA1 synapses, Schaffer collateral axons were stimulated using a twisted insulated nichrome wire electrode placed in CA1 stratum radiatum within 200– 300 μm of an aCSF-filled glass recording electrode, and paired-pulse facilitation (PPF) characteristic of this synapse (Wu and Saggau, 1994) was recorded. Baseline fEPSPs were obtained by delivering stimulus pulses with a 100 μs pulse duration applied at 0.1 Hz to generate fEPSPs between 0.3-0.4 mV in amplitude. Only experiments with ≤ 10% baseline variance were included in the final data sets. In addition, pairs of stimulation were delivered at a 50 millisecond (ms) inter-stimulus interval (ISIs). The paired pulse ratio (PPR) was calculated by dividing the initial slope of the second fEPSP by the initial slope of the first fEPSP.

#### 6.4 Long-Term Potentiation

At CA3-CA1 synapses, following a 20 min stable baseline (0.1 Hz, 100 μs pulse duration, with stimulation intensity set to elicit initial fEPSP amplitude of 0.3-0.4 mV), NMDA receptor (NMDAR)-dependent LTP was induced using high-frequency stimulation (HFS, 100 Hz, 0.5s s duration, repeated 5 times at a 20 s interval). The fEPSP slopes were normalized to baseline.

#### 6.5 Whole Cell Electrophysiology Recording in Acute Hippocampal Slices

For the recordings, slices were placed in a submersion recording chamber and perfused (3-4 mL/min) with aCSF. Experiments were performed at 28-32°C. For spontaneous EPSC (sEPSC) recordings, CA1 pyramidal cells were blindly patched on a Zeiss Examiner A1 upright microscope. Neurons were patched in the voltage-clamp configuration and recorded at a holding potential of −60 mV using a Multiclamp 700A amplifier (Molecular Devices). Patch electrodes (4–6 MΩ) were filled with a potassium gluconate based internal solution composed of the following (in mM): 130 K-gluconate, 0.1 EGTA, 20 KCl, 2 MgSO_4_·7H_2_O, 10 HEPES, 5 phosphocreatine-tris, 10 ATP, and 0.3 GTP. The pH was adjusted to 7.2 with KOH and osmolarity was 290-295. The access resistance and holding current (<200 pA) were monitored continuously. Recordings were rejected if either access resistance or holding current increased ≥ 25% during the experiment.

Analysis of sEPSC frequency and amplitude were performed using custom software written in Visual Basic, which measured amplitude and interevent interval. Events were fit to a template response and all events that fit the template and passed visual inspection were included in the analysis. The first 10 seconds of X-ray exposure was excluded from analysis as this was during the ramp up time of the Mini-X unit. Additionally, the 20 seconds after the Mini-X was turned off were excluded from analysis.

### 7. Intracellular cAMP Concentration Assay

Prior to performing the cAMP-Glo Assay (Promega, Cat. # V1502, Madison, WI) 20,000 HEK cells with stable expression of OptoXR-were suspended per well in a 96 well plate for treatment. A negative control group, not exposed to light during the duration of the experiment, established a baseline measurement of intracellular cAMP concentration. Exposures with either blue (470nm) LED light (5 seconds; 1mW/mm^2^) or X-ray in the presence of LSO:Ce particles (2 minutes) were conducted, followed by immediate lysis of the cells to preserve cAMP from degradation after stimulation. To ensure that X-ray emission alone was insufficient to induce intracellular cAMP, an X-ray exposure (2 minutes) without LSO:Ce RLPs was also performed. Relative light units (RLUs) were measured from the cell lysates following addition of a cAMP-dependent luciferase reagent in a Tecan Infinite F200 Microplate Reader. cAMP concentrations were calculated based on a cAMP standard curve according to manufacturer instructions. The sum of four technical replicates and three biological replicates are described here.

### 8. Materials

Commercial lutetium oxyorthosilicate particles (median particle size: 4 μm) were purchased from Phosphor Technologies and were doped with cerium at a 1-10 atomic % cerium (LSO:Ce). Prior to their use, the particles were washed with deionized water and vacuum air dried.

### 9. Statistics

All statistics were performed with Origin software (Origin Lab Corporation, 2002). Data are presented as mean ± standard error of mean (SEM). Samples sizes (n) refer to either cell or slice number for electrophysiological experiments. For most experiments, statistical comparisons were made using either student’s t-test or two-way ANOVA followed by a Tukey post hoc test. In the case of the cAMP concentration assay, a one-way ANOVA was used to determine a significance. Significance was determined as being p<0.05.

## Results

### Synaptic transmission is not affected by acute application of X-rays

First, we asked if basal synaptic transmission is altered by acute application of X-ray exposure. We recorded extracellular dendritic field excitatory postsynaptic potentials (fEPSPS) from CA1 pyramidal cells in stratum radiatum in response to CA3 Schaffer collaterals stimulation (Figure 1A). For each experiment, a stable 30 minute baseline was obtained and then the slices were exposed to either 30 minutes of X-rays (0.042 Gy/s) or unexposed (no X-rays) for the same amount of time. The initial slope of the fEPSP, representing glutamatergic postsynaptic response, was measured continuously throughout the experiments. We found no change in the fEPSP slope before or during X-ray exposure compared to unexposed controls (Figure 1B). In addition, X-ray exposure elicited no change in the paired-pulse ratio (PPR), an indirect measure of presynaptic glutamate release (Dobrunz and Stevens, 1997), (Figure 1C,D). Altogether, our data suggest that at this X-ray dosage (< 76 Gy) there are no significant effects on basal synaptic transmission.

**Figure 1.**
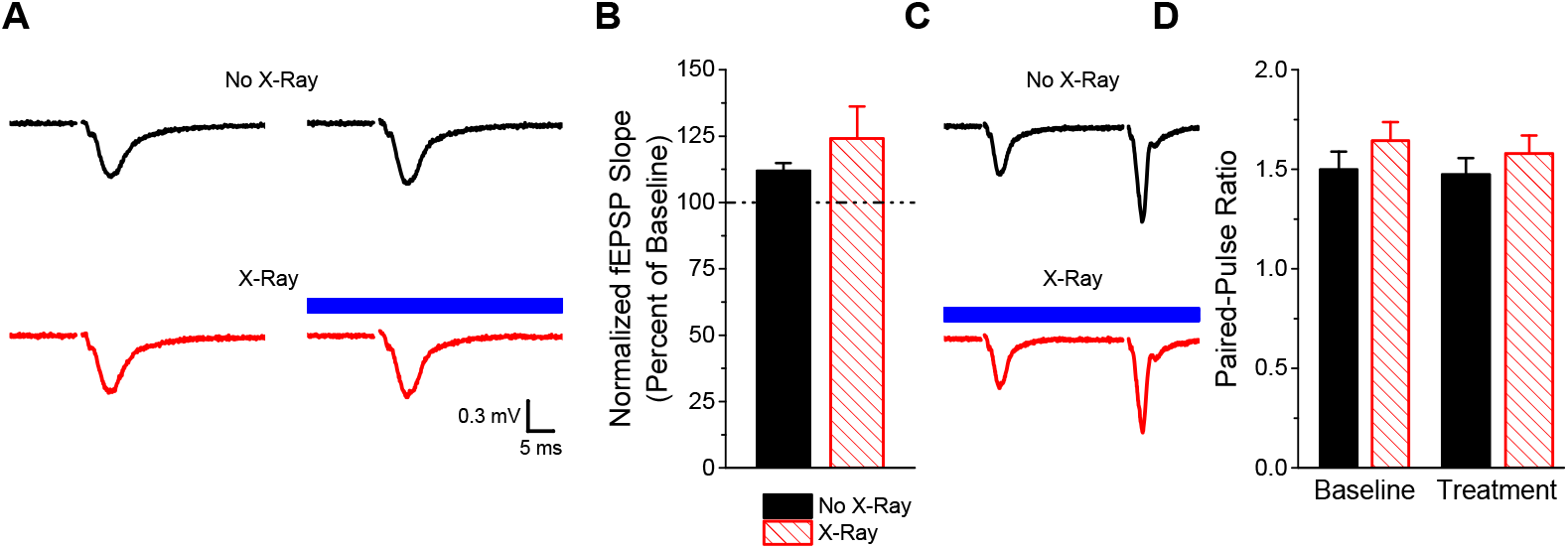
X-ray exposure has no effect on basal synaptic transmission. A) Example traces of fEPSPs in response to Schaffer collateral stimulation in acute hippocampal slices for the two conditions. The top traces (black) are from a slice that was never exposed to X-rays. The bottom traces (red) are from a slice that was exposed to X-rays for 30 mins after a stable baseline was obtained. The left trace is taken at the end of a 30 mins baseline period for each condition. The right trace is with (bottom) and without (top) X-ray exposure, taken 30 mins after the end of the baseline period. B) No change in the basal synaptic transmission with a 30 minute X-ray exposure (n = 7, 10; t-test; p > 0.05). C) Example fEPSPs traces of paired pulses at 50 ms interval with and without a 30 min X-ray exposure. D) No change in the paired-pulse ratio following a 30 min exposure to x-rays (n = 7, 10, 7, 10; ANOVA; F(_3,30_) = 0.046, p = 0.83).

### LTP is inducible after an acute, continuous exposure to X-rays

Next, we asked if long-term plasticity, a mechanism required for normal learning and memory, is compromised by 30 minute, continuous exposure to X-rays. Long term potentiation (LTP), an increase in synaptic strength that can last for hours *in vitro* to days when generated *in vivo*, has been extensively characterized at Schaffer collateral synapses onto CA1 pyramidal cells (Lømo, 2003; Nicoll, 2017). We recorded extracellular fEPSPs in response to Schaffer collateral stimulation in an interleaved fashion where experiments from X-ray exposed and unexposed slices were alternated. Following establishment of a 30-minute baseline of recording fEPSPs, slices were exposed to either 30 minutes of X-rays (0.042 Gy/s) or no no X-rays, followed by delivery of a high frequency stimulation (HFS, 100 Hz for 0.5 seconds, 5 times, separated by intervals of 20 seconds) to the Schaffer collaterals to induce LTP at CA3-CA1 synapses. This protocol produced identical post-tetanic potentiation and LTP magnitude in both X-ray exposed and unexposed slices. Indeed, no significant difference in LTP expression at 20 or 40 min post-tetanus in the X-ray exposed versus control slices was observed (Figure 2). Our data show that LTP is still inducible after an acute, continuous exposure to x-rays.

**Figure 2.**
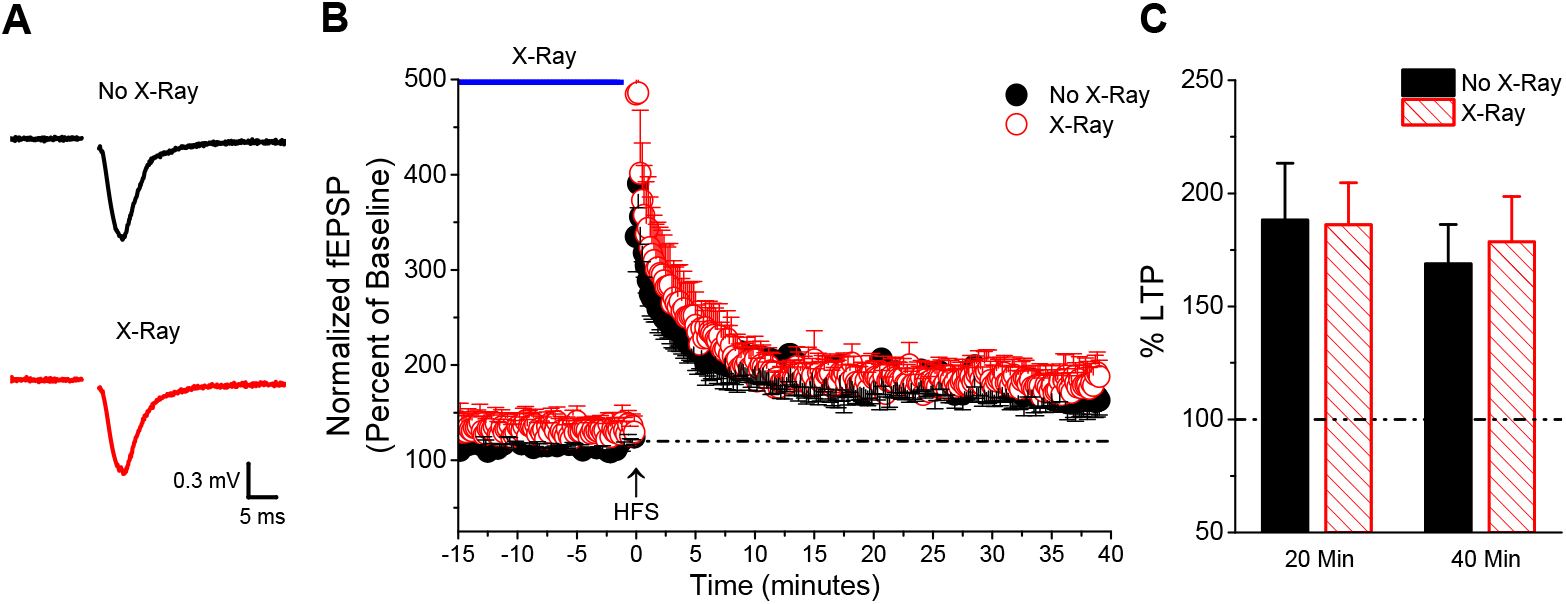
Prolonged, continuous X-ray does not prevent the induction of LTP. A) Averaged representative fEPSP trace at 35 minutes post-HFS stimulation with and without 30 min X-ray exposure. B) Induction of HFS LTP was similar between groups (n = 7, 9; t-test; p > 0.05) C) No difference in the LTP magnitude at either 20 or 40 minutes after induction with X-ray exposure (n = 7, 9; t-test; p > 0.05).

### X-ray exposure does not directly activate ChR2

Several photosensitive proteins can be activated by X-rays exposure (Dawson and Wiederwohl, 1965; Pande et al., 2016), especially ones that respond to UV light (Cannon et al., 2019). Channelrhodopsin-2 (ChR2), a light-gated cation channel, can be activated by wavelengths in the UV range (Bartley et al., 2019), as well as visible light (Boyden et al., 2005). However, it is unknown if ChR2 can be directly activated by acute X-ray exposure. To investigate this, we used Emx:ChR2 mice, in which ChR2 is expressed in excitatory pyramidal cells, to determine if ChR2 can be directly activated by X-rays. Whole-cell voltage clamp recordings of CA1 pyramidal cells were obtained, and the holding current was measured with and without X-ray application (2 min each, 3 cycles, 15 Gy, Figure 3A). Activating ChR2 allows the influx of positive ions that would depolarize the cell and force the amount of current needed to maintain the neuron at −60 mV to change, referred to as the photocurrent. No visible photocurrent was observed during application of 0.042 Gy/s X-rays, the rate of exposure performed in these experiments (Figure 3B). Additionally, this was verified by there not being a significant effect on the holding current in ChR2 expressing neurons (Figure 3C). This confirms that this X-ray dose does not directly activate ChR2 in the soma where the recordings are occurring.

**Figure 3.**
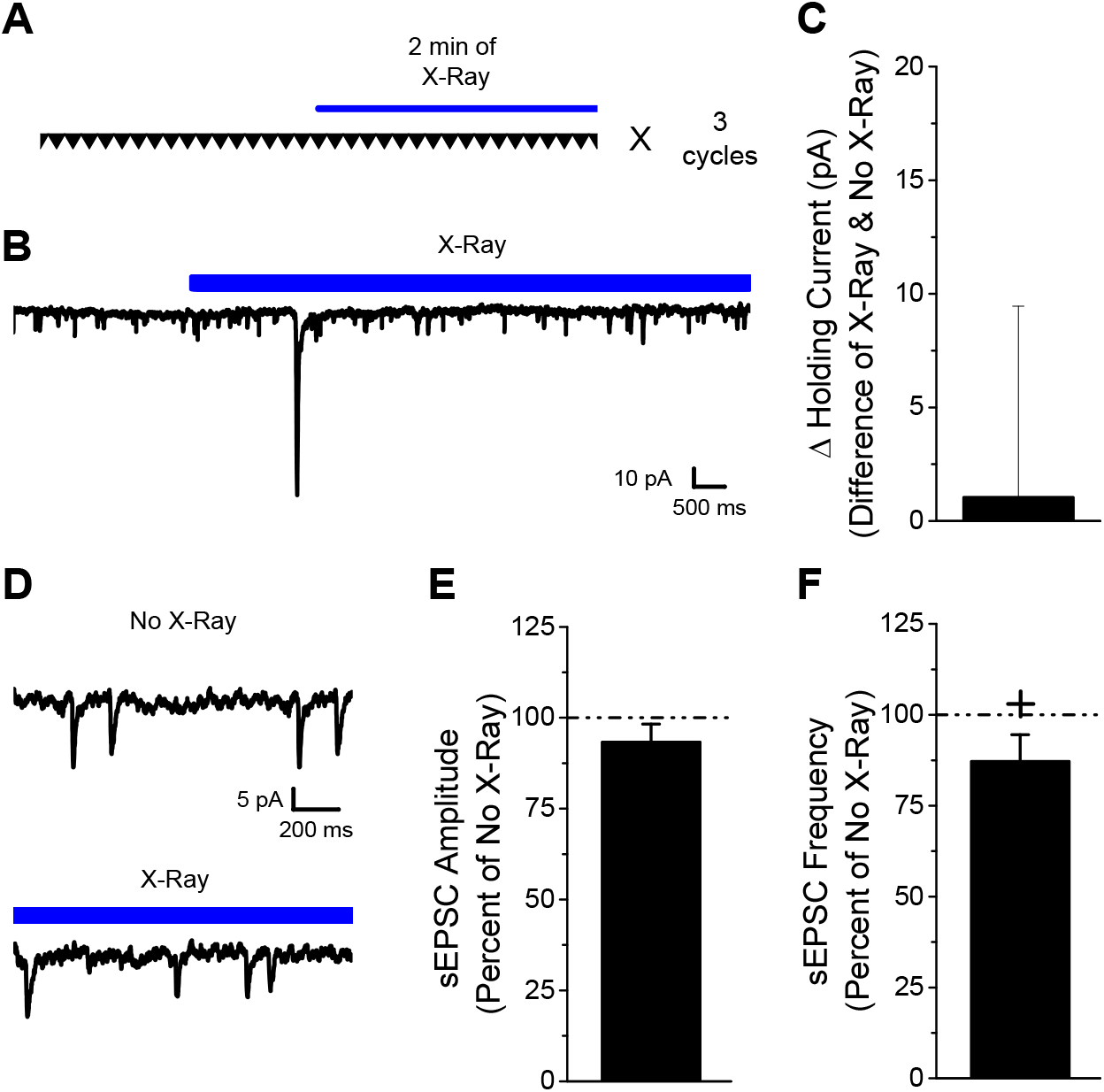
X-ray exposure does not directly activate ChR2. A) Schematic of experimental paradigm. B) Example traces showing that there was no photocurrent induced by X-ray exposure in neurons that express ChR2 C) No alteration in the amount of holding current needed to keep the PC at −60 mV (n=6; t-test; p=0.6). D) Example traces of sEPSCs onto CA1 PCs from Emx:ChR2 slices in the presence or absence of X-rays. E) No change in the sEPSC amplitude with X-ray exposure (n=6; t-test; p = 0.3). F) Minor decrease in the number of sEPSC events in the presence of X-rays (n = 6; t-test; p = 0.005).

However, X-rays might cause activation of ChR2 in either axons or dendrites thereby altering synaptic transmission. Therefore, we analyzed spontaneous excitatory postsynaptic currents (sEPSCs) onto ChR2 expressing neurons. As there is cell to cell variability with sEPSCs, we used a within cell control (no X-ray) to normalize the amplitude and frequency of sEPSCs. Figure 3A depicts the experimental paradigm showing that the periods of X-ray exposure and no exposure were interleaved throughout the course of the experiment. No change in the amplitude of sEPSCs was observed, but there was a minor decrease in the frequency of sEPSCs with the application of X-rays. Together, the data strongly supports that X-rays are not able to directly activate ChR2.

### LSO:Ce radioluminescence is appropriate for ChR2

ChR2 has been shown to be activated by wavelengths between 350 to 525 nm, with a peak at 470 nm (Lin et al., 2009; Bartley et al., 2019). Previously, we demonstrated that radioluminescent LSO:Ce microparticles are biocompatible with neuronal health and function (Bartley et al., 2019), and thus potentially useful for X-ray optogenetics. LSO:Ce has been widely studied as a scintillator, and has been shown to emit visible light in response to X-ray (Spurrier et al., 2007; Valais et al., 2008; Burdette et al., 2019). The LSO host lattice provides two potential Lu^3+^ sites that Ce^3+^ can substitute, with the Ce1 site surrounded by 7 oxygen neighbors while the Ce2 site has 6 oxygen neighbors. The relaxation of electrons from the 5d to 4f states of cerium from these two sites creates a broad radioluminescence emission that spans between 370-590 nm (Wu et al., 2019). We confirmed the emission spectrum for the 4 μm LSO:Ce microparticles in response to X-rays from the mini-X used in our experiments with a fiber bundle connected to a MicroHR monochromator. Figure 4 shows that LSO:Ce particles emit visible light in response to X-rays over wavelengths from 370-590 nm, which is well within the appropriate range to activate ChR2.

**Figure 4.**
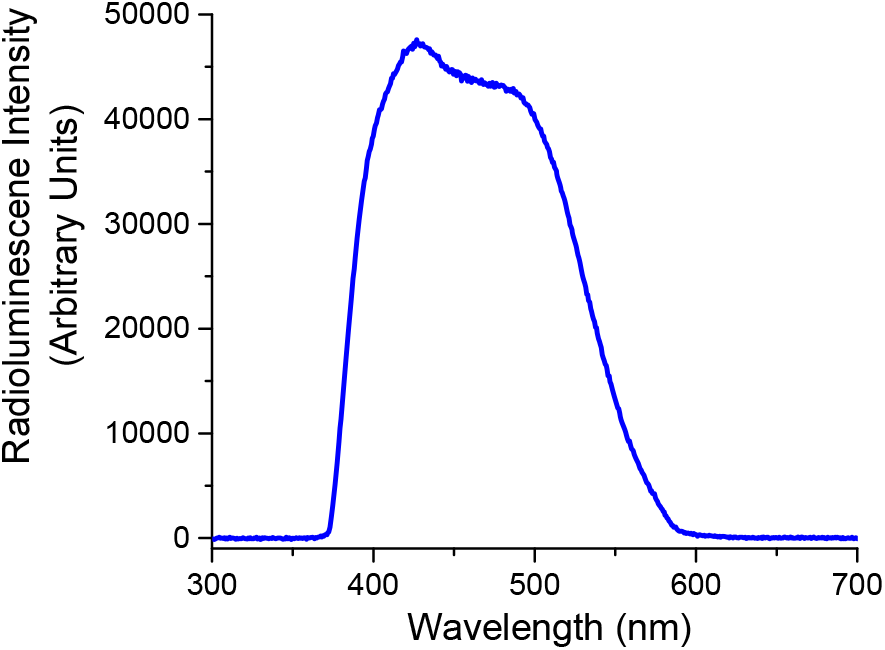
LSO:Ce microparticle emission is in range to activate ChR2. Emission spectrum of LSO:Ce microparticles in response to X-ray exposure. The peak emission of LSO:Ce microparticles is approximately 420 nm. ChR2 peak activation wavelength is around 450 nm.

### X-ray induced LSO:Ce radioluminescence stimulates G-protein coupled receptor activity

We first tested *in vitro* whether LSO:Ce RLPs could generate sufficient light output in response to X-rays to regulate opsin activity using a chimeric opsin receptor strategy. Of this class of chimeric receptors, collectively called OptoXRs, we utilized a chimeric fusion protein made up of light-gated extracellular ChR2 domain fused to the intracellular portions of β2-adrenergic receptor (β2-AR), and tagged with a fluorescent EYFP molecule on the C terminus to allow for easy visualization of transgene expression (OptoXR-EYFP). In response to light activation, OptoXR-EYFP undergoes a conformational change to trigger activation of G_αs_ signal transduction and stimulates cyclic AMP (cAMP) production mediated by adenylyl cyclase activation (Airan et al., 2009; Kushibiki et al., 2014). For these experiments, we expressed OptoXR-EYFP in HEK293T cells (Figure 5A). We then tested whether intracellular cAMP concentrations could be increased under various stimulation conditions using a cAMP-dependent luciferase reaction as a readout (see methods). As a positive control, we used LED stimulation in the absence of LSO:Ce RLPs, which caused a significant increase in cAMP concentration ([cAMP]) compared to cells that remained in dark conditions (Figure 5B). In order to determine whether LSO:Ce radioluminescence could also increase [cAMP], we exposed HEK293T cells expressing OptoXR-EYFP to X-rays in the absence or presence of LSO:Ce RLPs. Importantly, while X-rays alone did not alter intracellular [cAMP] relative to dark control, X-ray exposure in the presence of LSO:Ce particles caused a significant increase in [cAMP] (Figure 5B). Since cAMP is the primary downstream target of β2-adrenergic receptor activity, an enhancement in cAMP levels in the presence of X-rays and RLPs demonstrates that light emitted from the RLPs is capable of activating a chimeric opsin protein and causing downstream activation of an intracellular second messenger cascade.

**Figure 5.**
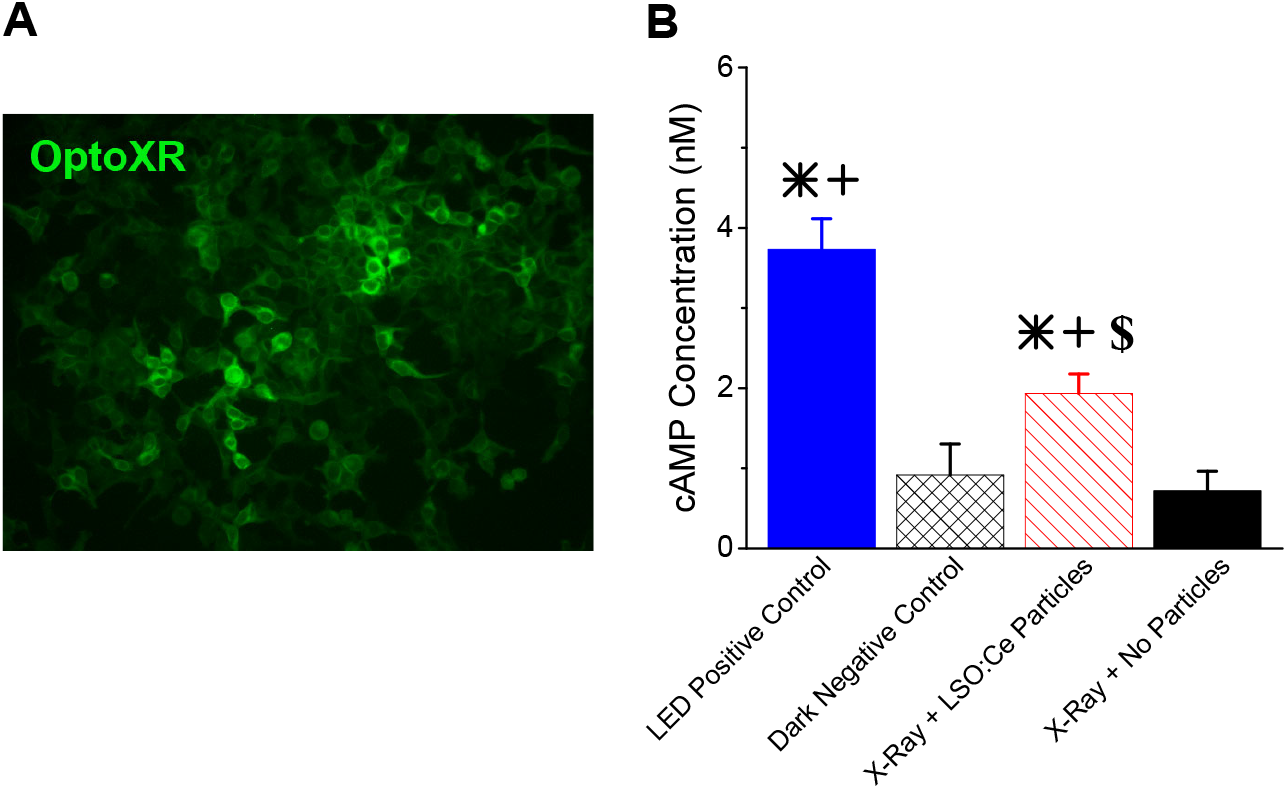
Radioluminescence from LSO:Ce particles induces cAMP production through light-gated β2-AR receptor (OptoXR) in vitro. A) Epifluorescent image of HEK293T cells stably expressing OptoXR-EYFP. B) The greatest intracellular cAMP concentration was measured in the LED control group, while the X-ray with RLP condition showed greater cAMP levels than either negative control conditions (n = 9/group; ANOVA; F_(3,32)_ = 29.1, p < 0.0001). * indicates significant difference from dark negative control (p<0.05). + indicates significant difference from X-ray + no particles (p<0.05). $ indicates significant difference from LED positive control (p<0.05).

### X-ray induced LSO:Ce radioluminescence stimulates neuronal activity

To determine whether X-rays can alter neuronal network activity via RLP-mediated scintillation, we recorded from dissociated hippocampal neurons expressing ChR2-mCherry between 16-18 days in vitro (DIV) that were either exposed to LSO:Ce RLPs or were not treated with RLPs. We recorded from neurons lacking expression of ChR2-mCherry and measured spontaneous excitatory post synaptic currents (sEPSCs) in the absence of the sodium channel blocker tetrodotoxin (TTX). These neurons received alternating 30s periods without and with X-ray exposure which provided less than 2 Gy of X-rays to each cell (Figure 6A). Figure 6B shows example traces of sEPSCs from neurons incubated with and without RLPs during periods of recordings in the presence or absence of X-Rays. Figure 6C displays pooled data for mean amplitude and frequency of sEPSCs for these conditions. Data are plotted as the ratio of recorded activity during X-ray exposure to activity recorded in the absence of X-ray exposure (mean of 3 repetitions for each condition). Interestingly, only neurons that received X-ray exposure in the presence of LSO:Ce RLPs displayed a significant increase in sEPSC frequency, while neither X-rays alone or RLPs in the absence of X-ray expsoure showed differences in sEPSC frequency or amplitude (Figure 6C-D).

**Figure 6.**
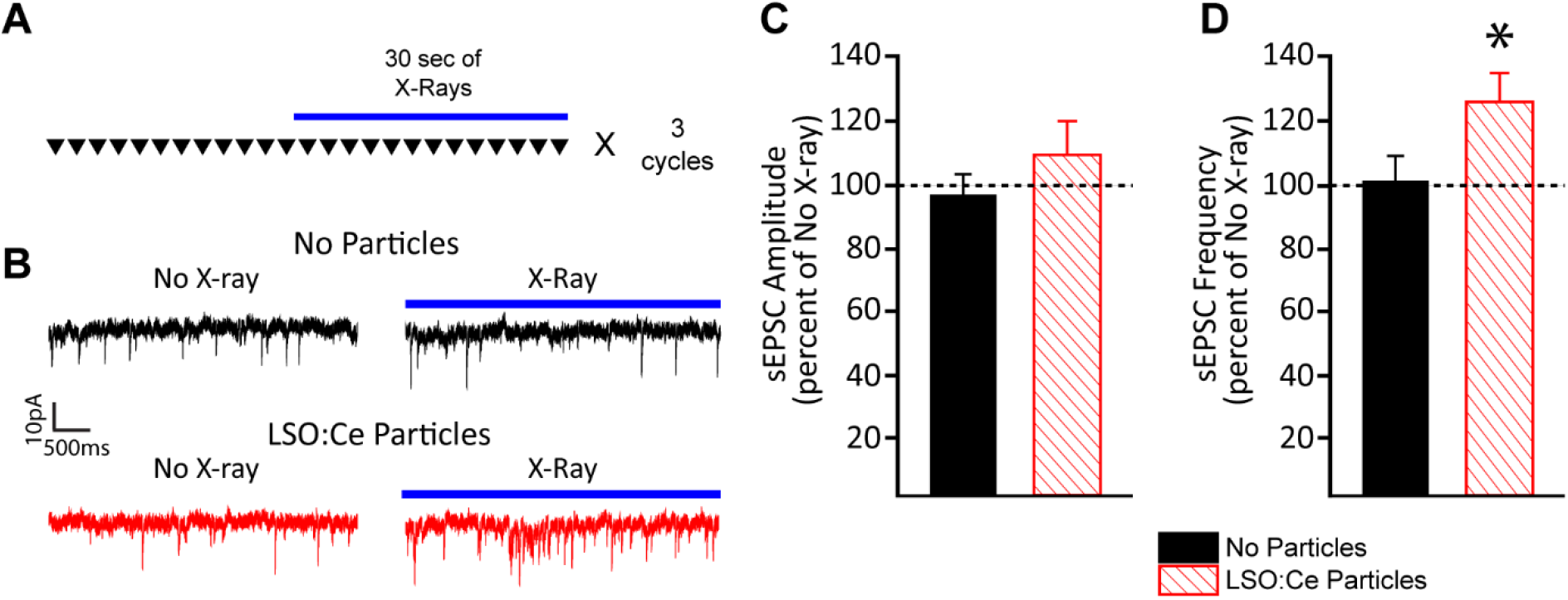
X-ray mediated radioluminescence from LSO:Ce particles enhances synaptic activity in primary hippocampal neuronal cultures. A) Diagrammatic summary of experimental design using alternating exposure to X-rays or absence of X-rays for each recorded cell. B) Representative voltage-clamp traces of sEPSCs recorded from hippocampal neurons with and without LSO:Ce particles in the presence or absence of X-rays. C) Pooled data illustrating that 30 second X-ray exposure did not alter sEPSC amplitude either in the presence of LSO:Ce particles or in the absence of particles (n = 7/group; ANOVA; F_(3,24)_ = 0.39, p = 0.76). D) Pooled data illustrate that exposure of cultures to X-rays in the presence of the LSO:Ce particles enhanced sEPSC frequency (n = 7/group; ANOVA; F_(3,24)_ = 3.76, p = 0.02) compared to periods when X-ray was turned off (dashed line). However, X-ray exposure alone did not alter the number of sEPSC events in the absence of LSO:Ce particles. * indicates significant difference between with and without LSO:Ce particles.

Next, we confirmed that the increase in sEPSCs using the combination of RLPs and X-rays can occur in acute hippocampal slices. We used Emx:ChR2 mice and incubated the slices with either LSO:Ce particles or vehicle. CA1 pyramidal cells that express ChR2 were used for the recording. Figure 7A shows the experimental paradigm and that the neuron received alternating 2 minute periods with and without X-ray exposure, thereby exposing the slice to approximately 15 Gy of X-rays. Figure 7B shows example traces of the sEPSCs recorded from ChR2 expressing neurons for the various conditions. We observed a small increase in the amplitude of the sEPSCs in the presence of LSO:Ce particles and X-ray exposure (Figure 7C). Importantly, we also observed a significant increase in sEPSC frequency with the combination of RLPs and X-ray exposure (Figure 7D). These data indicate that the X-ray induced radioluminescence from LSO:Ce particles is able to activate ChR2 and enhance network effects, as indicated by the increase in sEPSCs frequency and amplitude.

**Figure 7.**
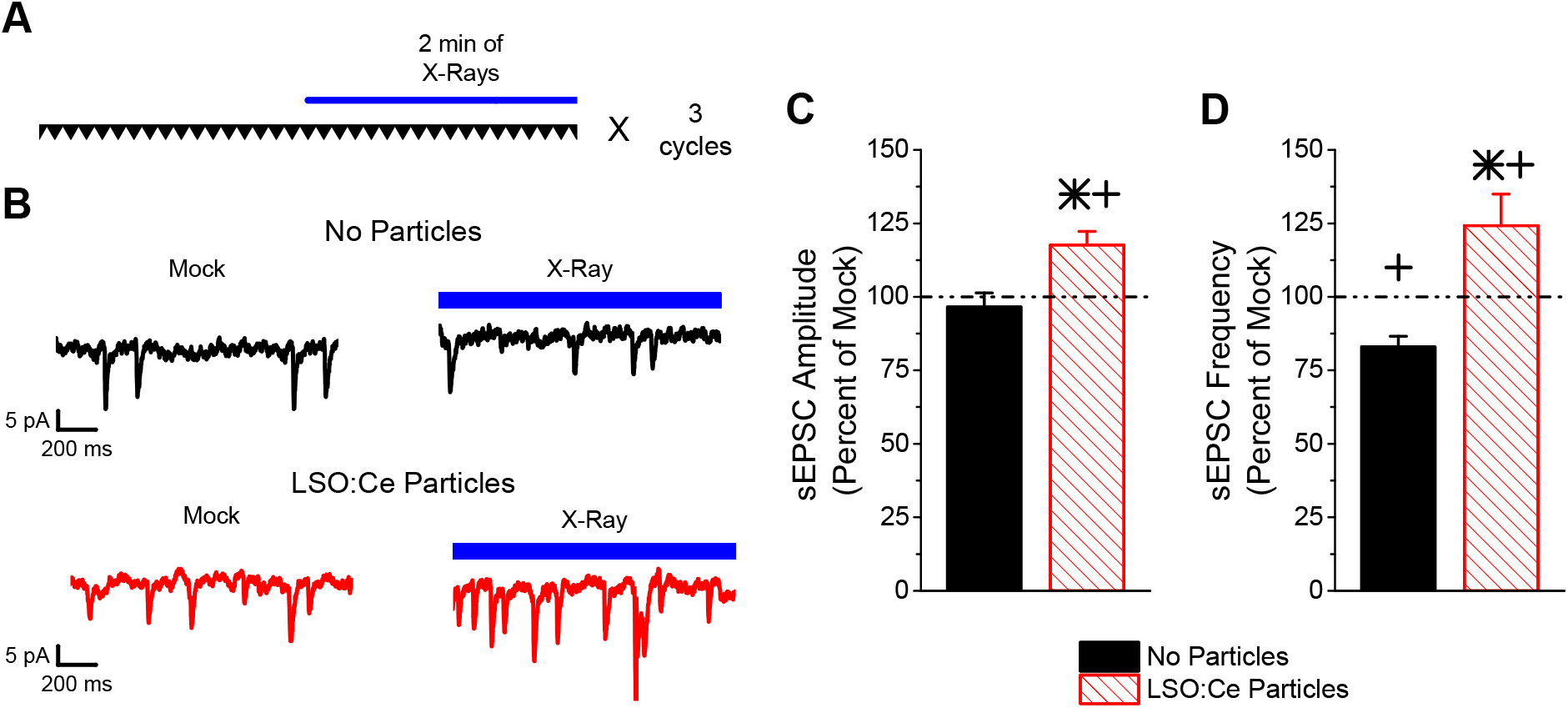
X-ray mediated radioluminescence from LSO:Ce particles enhances synaptic activity of ChR2-expressing neurons in acute hippocampal slices. A) Diagrammatic summary of experimental design using alternating exposure to X-rays or absence of X-rays for each recorded cell B) Example sEPSC traces onto CA1 PCs in slices from Emx:ChR2 mice, incubated with and without LSO:Ce particles, in the presence or absence of X-rays. C-D) X-rays plus LSO:Ce particles activated ChR2 in 4 out of 6 experiments. C) The presence of the LSO:Ce particles enhanced the sEPSC amplitude during X-ray exposure (n = 6, 4; ANOVA; F_(3,16)_ = 6.74, p = 0.004). D) The light emitted from LSO:Ce particles by X-ray activation increases the frequency of sEPSCs (n = 6, 4; ANOVA; F_(3,16)_ = 12.35, p = 0.0002). However, X-ray exposure in the absence of LSO:Ce particles reduced the frequency of sEPSCs recorded from PCs in Emx:ChR2 mice. * indicates significant difference between with and without LSO:Ce particles. + indicates significant difference between no X-ray and X-ray exposure.

## Discussion

Our results provide proof of principle that radioluminescence from scintillators can stimulate opsins and modulate synaptic function. Specifically, we show that X-ray induced visible light from LSO:Ce RPLs increased synaptic function via activation of ChR2. We also demonstrate that an acute 30 min X-ray exposure does not negatively impact hippocampal basal synaptic transmission or the induction of LTP. Furthermore, we showed that ChR2 is not directly activated by X-rays. Together, these results suggest that the combination of X-rays and LSO:Ce particles could be suitable for *in vivo* optogenetics using ChR2.

LSO:Ce is a scintillator that has been extensively characterized (Melcher and Schweitzer, 1992; Roy et al., 2013) and used in multiple medical devices (Stratos et al., 2010; Roy et al., 2013; Burdette et al., 2019). Cerium forms the luminescence center in LSO, and the emission spectrum makes it well suited for optogenetics applications using ChR2 (Berndt et al., 2011; Lin, 2011). We previously demonstrated that LSO:Ce microparticles are non-toxic to neurons and had very minor effects on synaptic transmission (Bartley et al., 2019). In this study, proof of principle experiments were performed showing that the combination of X-ray exposure and LSO:Ce RPLs are capable of increasing synaptic activity as indicated by enhanced sEPSC frequency in both cultured neurons and acute hippocampal slices. Additionally, we showed that the combination was able to work with a chimera opsin receptor and increase cAMP concentrations, showing some of the versatility of this system.

The use of X-rays has several distinct advantages over implanted LEDs to activate ChR2 in vivo. One disadvantage of implanted LEDs is that brain tissue causes light scattering, which therefore necessitates higher levels of light to activate ChR2 in vivo. This can cause heating of brain tissue; it has been reported that local tissue temperature increased by 0.8 °C with illumination used for optogenetics (Senova et al., 2017). Prolonged illumination and trains of light pulses elicit even higher temperature increases in tissue (Stujenske et al., 2015; Chen et al., 2018). Heating of tissue due to illumination causes damage and contributes to behavioral changes or physiological effects (Long and Fee, 2008; Stujenske et al., 2015). In contrast, application of X-rays for a short time caused no heating in the tissue (Matsubara et al., 2019). Another advantage of combining X-rays with RLPs is that scintillators will be extremely close to the cells expressing the opsins and therefore less light is needed to stimulate the opsin. Additionally, X-rays can penetrate further and retain the majority of their transmission compared to near infrared and visible light (Matsubara et al., 2019). Perhaps most importantly, X-rays penetrate the skull, removing the need for an invasive delivery method that causes physical damage.

Extremely high doses (>450 Gy) of X-ray have been shown to abolish nerve conduction (Gerstner et al., 1955; Gerstner, 1956) and alter synaptic transmission (Portela et al., 1977). Deficits in fear memory can occur as a result of acute X-ray exposure, independent of its effect on neurogenesis (Puspitasari et al., 2016). However, in our study, with a moderate dose of X-rays (< 80 Gy), no effect was seen on basal synaptic transmission and LTP could still be induced. LTP was even maintained to the same level even 40 minutes after induction. It is possible that waiting a longer time period after X-ray application might have different effects on synaptic transmission, as one study showed a transient decrease in dendritic protein levels hours after the exposure (Puspitasari et al., 2016). However, it is most unlikely that low doses (< 10 Gy) of X-ray will influence synaptic transmission and these are levels that can be used for optogenetics, as we have shown here. In our study, we show that using less than 2 Gy of X-rays to mediate scintillation from LSO:Ce particles is sufficient to cause changes in neuronal activity via ChR2.

Multiple proteins can be altered and/or activated by X-rays, including rhodopsin (Lipetz, 1955; Dawson and Wiederwohl, 1965; Groh et al., 1984; Fuglesang et al., 2006; Sutton et al., 2013). Recently, LITE-1, a UV light sensitive G-protein coupled receptor (GPCR) in C. elegans, has been discovered to also be activated by X-rays, producing behavioral effects (Cannon et al., 2019). However, to our knowledge, none of the opsins typically used for optogenetics have ever been tested against X-ray exposure. Several of these opsins can be activated by a wide range of wavelengths, including ChR2, which can be activated by UV as well as visible light (Bartley et al., 2019). We did not detect a photocurrent in response to X-ray stimulation in neurons that expressed ChR2. However, this was recorded at the soma and it is possible that a small photocurrent was induced by X-ray activation of ChR2 in the dendrites or axons. But, we also saw no increase in synaptic activity during the application of X-rays alone in neurons that express ChR2. Together, these results suggest that ChR2 is not directly activated by X-rays. Further studies will be needed to determine if this is true for other opsins used in optogenetics.

Repeated X-ray exposure has been shown to abolish neurogenesis (Burghardt et al., 2012; Lacefield et al., 2012) and effect cognitive function, as seen for radiotherapy (Pazzaglia et al., 2020). Neurogenesis can be diminished even with a single X-ray exposure (Mizumatsu et al., 2003; Casciati et al., 2016). However, this is most likely due to oxidative stress and can be minimized using a free radical scavenger (Motomura et al., 2010). Low doses of X-rays had minimal to no effect on neurogenesis (Casciati et al., 2016; Prager et al., 2016). In our studies, we did not analyze the effect on neurogenesis for the doses used, but our proof of principle experiments reported here employed doses that are in range with minimal effects on neurogenesis. A study has shown that obtaining the peak X-ray luminescence emission from scintillators can be fine-tuned and requires less energy (George et al., 2019), suggesting that activation of the LSO:Ce microparticles could occur at even lower X-ray doses.

Other studies have shown that light emitted from various materials can activate different opsins (Ronzitti et al., 2017; Chen et al., 2018; Lin et al., 2018; Matsubara et al., 2019; Miyazaki et al., 2019; Gong et al., 2020). One study has shown that the use of X-rays and a scintillator can be used to induce circuit changes through activation of an opsin (Matsubara et al., 2019). However, they implanted a large crystal, which was larger than a fiber optic, that caused damage to the surrounding tissue. Additionally, there was a similar increase in active microglia with implantation of the crystal as compared to a fiber optic. In contrast, we previously showed that the LSO:Ce microparticles had no effect on astrocytes (Bartley et al., 2019). LSO:Ce microparticles therefore have the advantage of being able to enhance synaptic transmission through X-ray induced luminescent activation of ChR2 while avoiding tissue damage that can occur with other materials.

Even though X-rays can transmit at a high penetrance for several millimeters, the X-ray output can be diminished if the target is too far from the source. In our experiments, the distance from the X-ray source to the cells was variable. This would cause some slight differences in the amount of X-irradiation that occurred for each experiment and probably contributed to the small effect seen in our proof of principle experiments. One of the limitations of our current experimental paradigm is that the particles were applied to the surface of the slice, and where therefore more than 100 microns from the cells expressing ChR2. In addition, the particles are not evenly dispersed across the slice or coverslip. This partially accounts for the modest effect of ChR2 activation observed in this study.

In summary, our results provide the first demonstration of enhanced neuronal activity by X-ray activation of radioluminescent microparticles. While further studies are needed to determine the appropriate size of LSO:Ce particles that can be used in vivo, as well as the necessary density required, our study supports the possibility of using radioluminescence from LSO-Ce particles caused by x-ray activation as a noninvasive way to deliver light to the brain.

## Notes

### Competing Interest Statement

The authors have declared no competing interest.

